# Frequency Transformations and Spectral Gaps in Cochlear Implant Processing

**DOI:** 10.1101/035824

**Authors:** Barry D. Jacobson

## Abstract

In previous work[1] we created a mathematical model and identified a major source of distortion in Cochlear Implant(CI) processing which manifests itself in three forms, all of which are due to the nonlinear envelope processing algorithms which are widely used in some form or another in many current models. The first are spectral gaps or dead zones within the claimed frequency coverage range. This means that there exist regions of the spectrum for which there is no possible input that can produce an output at those frequencies. The second are frequency transformations which convert input tones of one frequency to tones of another frequency. Because this is a many-to-one transformation, it renders following a melody impossible, as the fundamental frequency of two different notes may be mapped to the same output frequency. This makes them impossible to distinguish, (although there may be differences in higher order harmonics that we will discuss). The third type of distortion are intermodulation products between input tones which yield additional output tones that were not present in either input. In the case of multiple talkers, these will compound the comprehension difficulty, as not only are the original spectral components of each speaker transformed, but additional nonexistent components have been added into the mix. This accounts for the great difficulty of CI users in noise.

In this work, we extend our earlier work in three ways. First, we clarify our description of spectral gaps which a number of readers pointed out was unclear, in that it implied that certain input tones will produce no response at all. In fact, all input tones will produce a response, but in most cases, the output will be frequency-transformed to a different frequency which the CI is capable of producing. Second, we graphically illustrate the input/output frequency transformation, so that the reader can clearly see at a glance how each frequency is altered. The form of this transformation is a staircase over most of the usable range, meaning that for single, pure tones all frequencies in the passband of a particular channel are mapped to a single frequency—the center frequency of that channel. As frequency continues to increase, all frequencies in the passband of the next channel become mapped to the center frequency of that channel, and so on. The exception is in the low frequencies, for reasons that we discuss. Third, in our earlier work we analyzed the simple case of only two pure tones within a single channel. Here we extend to the more realistic case of mixtures of complex tones, such as musical notes or the vowels of speech which may each have multiple harmonics extending throughout much of the audible frequency range. We find that, as expected, the output components of a source within a single channel often clash (are dissonant) with each other, and with those output components of that source (higher harmonics) which fall within other channels. So that instead of there being a harmonic or integral relationship among the output spectral components of each source, these components are no longer related to each other harmonically as they were at the input, thus producing a dissonant and grating percept. Furthermore, in the case two or more complex tones, additional intermodulation components are produced that further distort the sound. All these assertions are derived from theoretical considerations, and also noted from the author’s own listening experience, and further confirmed from correspondence with other CI users.

## 1 Introduction

As we related in our earlier work,[1] I had been a long time user of hearing aids since a hearing loss developed at around age 4. I was fully postlingual, and enjoyed listening to music at that age. I would often play various musical records on my home turntable. My nursery-school teacher noticed that I would often go up to the classroom record player and turn up the volume unusually loud, so she recommended to my parents that I get my hearing tested. I was found to have a sensorineural loss that stabilized for many years at around 90 dB. I was fitted for hearing aids, and had a completely normal childhood and early adulthood. I never considered the hearing loss to be a handicap any more than I considered wearing eyeglasses to be a handicap, which I was prescribed at about age 12 for routine nearsightedness. I had numerous friends, participated in all sports activities, played musical instruments and participated in choirs. Throughout, many audiologists and laypeople have told me my speech sounds normal. I note all this to emphasize that I know very well how speech and music are supposed to sound.

Unfortunately, a sudden acoustic trauma due to an overly high setting on a new hearing aid damaged my hearing to the point where I could no longer derive much benefit from hearing aids. I was evaluated and found to be a candidate for CIs. I chose the Med-El unit primarily because of their deep low frequency coverage, and for their claim of encoding temporal information in the form of waveform fine structure, rather than merely envelope information, as other manufacturers do. I had also hoped to preserve residual hearing, as they claim to have a very soft electrode which is supposed to minimize insertion trauma. But despite being operated on by the world-renowned surgeon, Dr. Thomas Roland, all residual hearing in that ear seems to be gone.

Our purpose in this work and the earlier work is to make it clear that CI users experience significant distortion which greatly impacts speech reception and music recognition, and to suggest what improvements can be made to CI processors in order to correct this problem. While under ideal conditions, CIs do provide much benefit in hearing and understanding speech, nevertheless, the distortion renders the sound quality buzz-like, raspy and grating, which is annoying and fatiguing, in addition to making it more difficult to understand than if the speech were clear but presented at the same loudness level. For music, the distortion is even more detrimental, and can make even the most well-known melodies unrecognizable, so that by the time one realizes what was playing, the song is over. The reason is that speech is more robust, and the general location of the formant peaks is more important than the exact frequency of the fundamental. However, in music, frequency is critical, and a change of only 6% in frequency is enough to change one note into another. But aside from that, as we will see, the harmonics do not properly track the fundamental, thus ruining the periodicity of the waveform. This changes a musical percept into a noise-like percept which makes recognizing pitch even more difficult. We emphasize that loudness is not the problem with CIs, as sounds are louder than I could achieve with hearing aids (the quest for loudness was what ruined my hearing in the first place); the distortion is the main issue that needs to be addressed.

We will make a brash and opinionated statement at this point to distinguish this work from what is common in the CI and hearing field. Most studies involve playing a particular stimulus to a group of CI users and trying to measure how close their response comes to that of normal hearing listeners or perhaps to users of other CI models, or CI programs, or hearing aid wearers. One may test a particular vowel, or a particular word or sentence set. In music one might test a particular note or melody. The subjects might be restricted to children, or new users, or experienced users, etc. One might further divide prelingual from postlingual subjects, or bilaterally impaired from unilaterally impaired, or one age range from another, and so forth. If a subject has bilateral impairment, one might compare a CI in one ear vs. a hearing aid in another. There are many combinations of subjects and test materials that can be tried. Each of these possible studies can yield its own research publication. Much use of statistics is made to quantify the performance on each task and with each group of subjects.

But our approach is different. We look at the copious amount of already-existing data, and see that in study A, 56% of subjects got correct answers on test set X, in study B 67% were correct on test set Y, and so forth. The only statistic which is of importance to us is that in all studies we would like to see 100% of users get 100% correct on 100% of the material. Anything less is a failure. We hear the distortion ourselves, and immediately understand the difficulty users experience on these tests. The exact numbers are not important to us. Our goal is to solve the problem once and for all. And the very first step is to understand what is going wrong, and what the source of the distortion is. The next step is for manufacturers and the hearing community to correct the problem through the design of more intelligent algorithms, so that performance is improved. Mere generation of endless statistics does not necessarily do a service to the hearing impaired community.

We have received objections that as an individual, a study where the number of subjects N is equal to 1 is not reliable and objective. Our response is very direct. We not only hear the distortion, but we also demonstrate mathematically what its source is, based upon publicly available information and block diagrams of commonly employed CI processing algorithms. We need to make only one solitary assumption to be able to translate this information into a workable and testable model, as we will discuss. Because nobody can imagine what electrical pulses sound like, we must rely on conceptually translating the output of such block diagrams into a form that can be analyzed using conventional signal processing concepts and building blocks. We allow that there is room for disagreement with how this should be implemented, but there are limits to what can be changed even if one wishes to substitute other alternative schemes. We further note that CIs, in general, follow a vocoder-inspired design, and our model treats the CI as being an exact analog of a vocoder. Thus, we believe the steps we have chosen are entirely appropriate to analyze the spectral response of a CI. We correlate as well as we can via careful listening the percepts we hear with the percepts that would be heard based upon the output of our processing model, and do not note any differences that would indicate that our model is incorrect. We describe this in detail later.

The advantage of using a model is that it gives a starting point and a framework for discussion as to what users are hearing. It also lays one’s cards out on the table, so that everything is transparent, and all assumptions are there to see. If a user disagrees with a step, which he is free to do, he must point to the exact place in the model which he feels is incorrect, and create a replacement. We have been frustrated in that some people who disagree with our views have tended to talk around our assertions in vague terms, but without specifying what exactly the alternative should be, and what effect it would have on the actual output. A model insures precision and focuses one’s thinking, and also eliminates the need for subjective descriptions. It is this author’s opinion that a major impediment towards progress in CI designs is the inability for users and engineers to communicate. Engineers cannot hear the sound directly, like they can see the picture of a TV they are designing. They must rely on users. But unfortunately, most users are not technically trained and can only use the most vague terms like “not clear”, or “noisy”, etc. But what the actual defect in the signal is, could be anyone’s guess. And does “noisy” refer to additive noise, or to the signal of interest, itself. So this lack of common language between users and engineers has made it very difficult to diagnose problems. And an incorrect diagnosis leads to incorrect design changes, causing possibly worse problems than were originally present.

Since this author has been trained in electrical engineering and signal processing, and is also a CI user, and is furthermore, a very fussy individual who listens extremely closely, and is not satisfied with anything less than 100% accuracy, it might be worth considering what he has to say.

Because the following discussion is relevant to everything we will be doing, we discuss the philosophy which led us to choose the model that we did, and the reasons for objections from at least one member of the hearing community, our close friend Dr. Aryeh Litvak, whom we anticipated in our earlier work would disagree. There is a major controversy raging in the hearing community over the relative importance of spatial vs. temporal cues regarding the encoding of frequency information in normal cochlear processing. On one hand, frequency appears to be represented in terms of the specific place along the basilar membrane of the cochlea that is maximally responsive to sounds of a particular frequency. Higher frequency sounds resonate at locations closer to the base of the cochlea, while lower frequency sounds resonate at locations closer to the apex. But on the other hand, frequency also appears to be encoded in terms of timing information of neural spike trains, with higher frequencies firing closer together in time than lower frequencies. This neural synchrony with waveform features such as local maxima, is known as phase-locking. The model we have chosen places more importance on temporal information than on spatial information, in that the ultimate determination of frequency must be consistent with the temporal features. For example, if a higher frequency tone is introduced at a slightly lower frequency place in the cochlea by means of electrical stimulation, our working assumption is that the tone will still register as a higher frequency tone. We believe that spatial information is important, but is used in combination with temporal information to simultaneously produce both the enhanced frequency resolution and enhanced temporal resolution that are necessary to decode speech. Were speech to be encoded in only a single place or channel within the cochlea, it would take a longer time to accurately measure its frequency. This is due to the uncertainty principle of Fourier Analysis, which states that to get higher temporal resolution, one must trade off frequency resolution, and vice versa. However, speech requires both, since syllables last only a short time, necessitating fast temporal resolution, but to decode them and separate from competing sounds, requires accurate frequency resolution, as well. In our Ph.D. thesis,[2] we showed that by combining information from multiple channels, each with a slightly different frequency response, one can meet both goals and separate closely spaced tones in frequency, but in a relatively short time.

Because we believe that, ultimately, temporal cues are most or at least crucially important for determining frequency, we can use standard signal processing techniques, such as analyzing the spectral content of an output waveform using Fourier methods, and comparing it to the input. However, if one follows the school of thought that spatial position actually determines frequency, regardless of the exact temporal waveform, one is at somewhat of a loss to distinguish the properties of two signals from each other, each introduced at the same cochlear location by electrical stimulation, as one would conclude that regardless, the output frequency would be the same. This latter school of thought clearly must place a very heavy emphasis on the importance of the placement of the electrode array within the cochlea, as its adherents believe that any slight shift in location will alter the frequency content up or down. However, according to the first viewpoint, such a shift might not be as significant, as ultimately the waveform shape will determine what the user hears.

Litvak and indeed much of the CI research community believe that spatial resolution is key. They base this on experiments which tried to measure frequency discrimination in CI patients, and concluded that not more than 300 Hz variation can be detected when electrical stimulation is introduced at a particular cochlear location. This number, 300 Hz, figures prominently in the CI design strategy, as we will see later.

Due to this belief in spatial primacy, the Advanced Bionics strategy is to attempt to increase the number of possible tone percepts by steering current into intermediate locations between electrodes. This is accomplished by stimulating two adjacent electrodes simultaneously in various relative strengths to focus current at specific points in the cochlea located in between the two electrodes. For example, if one wanted to reach a point halfway in between, one could apply equal strength pulses to the two electrodes. But if one wanted to reach a point closer to one electrode, then one would increase the strength of the pulse at that electrode, and correspondingly reduce the strength of the pulse at the other electrode. However, if temporal primacy holds, this strategy will not significantly improve frequency discrimination, as the exact place doesn’t matter, rather the temporal information matters, and that has for the most part been discarded by the envelope processing algorithms, as we will see.

Another major difference in approach between these two opposing views is the relative importance of the characteristics of the signal used to stimulate the electrodes. If one holds of spatial primacy, then the existence of distortion components will not be as relevant, as the spatial location will determine the frequency percept, regardless of the exact waveshape. Perhaps for that reason, nobody has bothered to analyze these components, to the best of our knowledge. In our opinion, this is the major misunderstanding that plagues CI design today. But the problem is actually compounded by a second misunderstanding. Even those researchers who do place importance on temporal information seem to believe that envelope processing is satisfactory, but adding temporal information might possibly enhance frequency resolution, perhaps making the percept less ambiguous. But in fact, the problem with envelope processing is not that it is ambiguous, but that it completely obliterates and rewrites the frequency content of the original signal. The symptoms are roughness, tonal distortion, missing frequencies and creation of spurious components. While we demonstrated this in our earlier work for single channels, the key point of this paper is to extend the analysis to all channels, and examine how they interact. We first plot the input/output frequency transformation across the entire response range of a CI for a single, pure-tone input. This will visually clarify a point which confused certain readers of our first work, regarding the existence of spectral gaps or dead zones. Second, we extend to complex tones by comparing the full spectral content of a vowel or musical note at the input and output. And third, we examine what happens with a mixture of two such complex tones, such as when two or more speakers are talking at once. We will find that intermodulation distortion components are created that make an already difficult situation for CI users even worse in such noisy environments.

Having chosen a model, we compute mathematically and illustrate graphically the exact output components that it predicts will be produced, and compare with the components present at the input. Significant differences are noted. We then relate these mathematical differences to the percepts one would be expected to hear. Finally, we suggest improvements in future designs to solve these problems

We note that in many situations, a group of subjects where *N* = 1 is sufficient to make a determination as to a course of action. If one’s car doesn’t start, one doesn’t convene a panel of 25 people to try to start the car, record how many successes they had, and compile exact statistics. One instead proceeds immediately to the mechanic based upon a single, solitary test subject, namely, the owner. And conversely, when Edison was inventing the light bulb and succeeded in getting one to burn long enough to be useful, he, too, didn’t need to convene a panel of subjects in order to gather statistics as to whether or not it worked. He could see for himself. And the same holds true for the invention of the telephone, which history records as there being only two people present, Alexander Graham Bell and his assistant. One spoke, one listened and they knew they had succeeded. A single user can ascertain whether a device works or not. It is sometimes wise not to complicate things more than necessary. Perhaps engineers think differently than scientists in that regard.

We also note that certain notions which are popular in the field may actually be very counterproductive. The idea that there exists some kind of magical auditory plasticity which will solve all our engineering problems and change distortion into clear sound is, in the opinion of this author, a fantasy. As we noted in our earlier paper, an incorrect eyeglass prescription doesn’t correct itself, nor do smeared lenses on a pair of glasses. A rolling or snowy television picture doesn’t become more viewable over time. The brain expects a correct input before it goes to work with its analysis. It may take a few days or a week to get used to the sound, but that is in the best case, where the output is as it is supposed to be. If certain components are missing, or nonexistent components have been added, or if certain frequencies have been transformed, there is little the brain can do, except try to decipher the incomplete message the best it can. This causes much fatigue, just as when l_t__rs are missing from a printed page.

Finally, we emphasize a point which we also noted in our earlier work, that our analysis indicates that the distortion is a result of faulty algorithms in the processor itself. It is not a result of interelectrode interference or current spreading, as commonly believed. It is not a biological phenomenon, but strictly an electrical engineering problem. This has major ramifications, as it calls into question the entire rationale for Continuous Interleaved Sampling (CIS) schemes and their variants. If we are correct, there is no reason to use interleaved or pulsed schemes to avoid interference that might occur between channels were pulses fired simultaneously, as that is not the source of the distortion. And furthermore, as we noted there, these introduce switching noise, that actually adds to the distortion. (This is a completely separate type of distortion, being an additive crackling and hissing noise, not a frequency transformation of the input, as is the other, primary, type of distortion we have been addressing.) In addition, if we are correct about the true source of the distortion, it also calls into question the issue of why Med-El, in particular, which has the longest electrode (31.5 mm) and the most space to work with, chose to limit to 12 channels, the fewest of all manufacturers, due to fear of such current leakage. (Other manufacturers use a shorter electrode and have more channels, and hence a closer physical spacing.) Our analysis shows that this fear is unfounded, as the actual source of the distortion is the nonlinear envelope processing algorithm. We will see that it can generate phantom frequencies that might have appeared to come from a different channel. Perhaps when users were interviewed, they described such interference. We surmise that this is what misled engineers into thinking channels were leaking from one to another. In truth, we believe that many more channels can be accommodated, with no reason to limit spatially due to fear of interference.

The conclusions of this work and our previous work are that many of the problems in speech intelligibility and music recognition that CI users experience are due to distortion that has been introduced due to an incorrect understanding of the input/output frequency relationships of current CI processing schemes. But the good news is that these appear correctable, and that furthermore, greatly enhanced frequency resolution should eventually be possible, as there is no biological limitation to the number of electrode channels that can be used; the only limitations being practical engineering considerations, such as wiring size, transmission through skin and power consumption.

## 2 Model and Analysis

A typical diagram of a basic CI processing scheme is shown in Figure 1. The signal is separated into frequency bands by a bandpass filter. It is then rectified to remove the negative-going parts of the waveform. This leaves only the positive half, and can be integrated, for example with a capacitor, and the value of the integral will be equal to the average amplitude in the integration interval. Were the signal not rectified, the integral would be zero, or close to it, as the positive and negative portions would cancel. It is then passed through a low-pass filter, which acts as an integrator, and also smooths or averages the signal to removes high frequency fluctuations and unpleasant discontinuities introduced by the nonlinear rectification process. The output of this step is effectively a moving average of the signal amplitude, which is known as the envelope. It tracks the signal strength over a time window which is approximately the reciprocal of the cutoff frequency of the lowpass filter.

**Figure 1.**
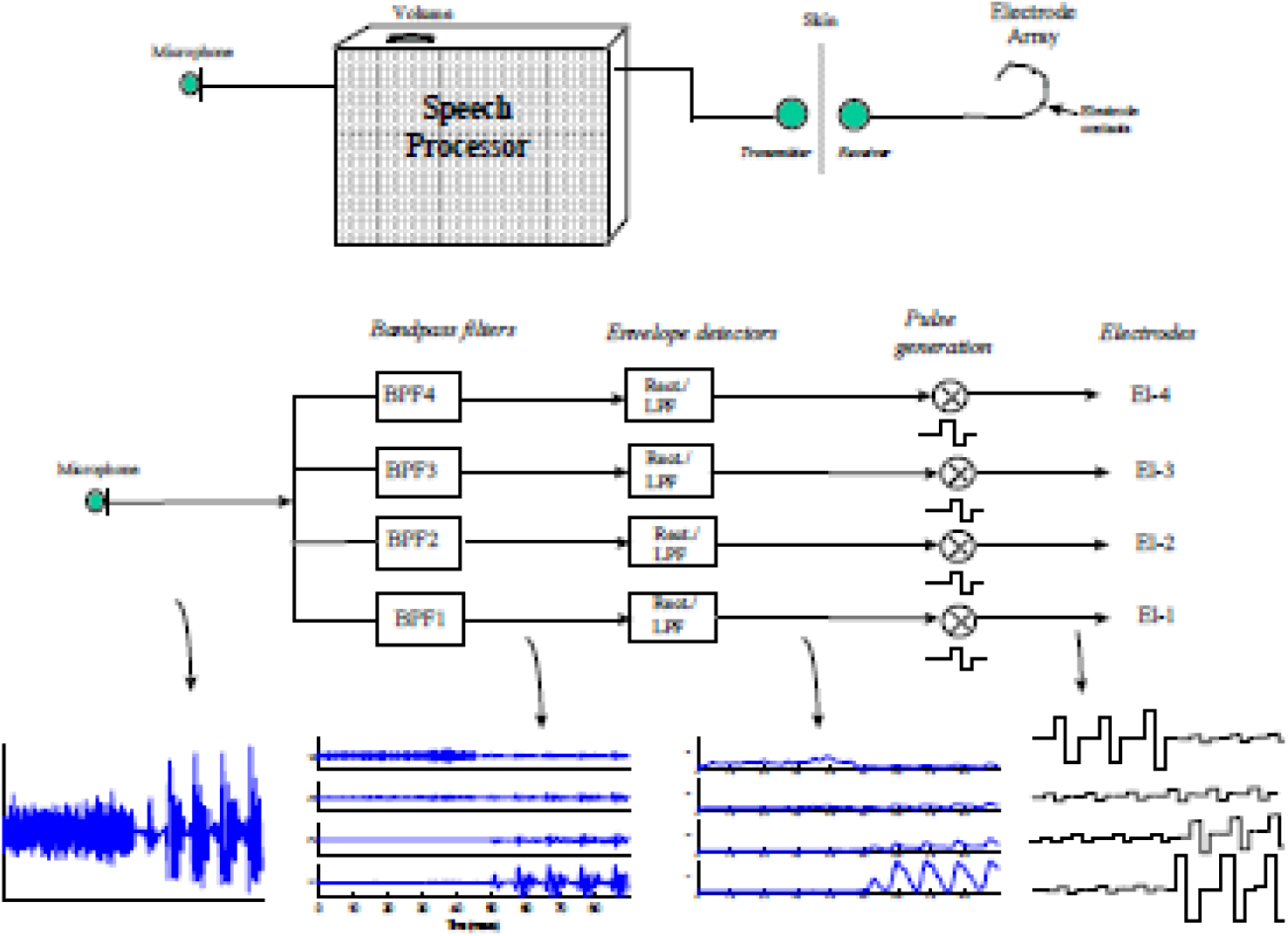
An illustration of a typical CI processing scheme. The signal is passed through a filter bank, and then each channel is rectified and low-pass filtered. The resulting envelope is used to modulate a pulse train. Note that in a Continuous Interleaved Sampling (CIS) scheme, the timing of the pulses in each channel would rotate, so that no two channels would fire at the same time. From Loizou.[3]

The final step in an actual CI, is to use the envelope to modulate a pulse train which directly stimulates the auditory nerve in accordance with the average signal strength of the time period. But here is where we must substitute an alternate step, as we indicated in the Introduction, if we are to analyse the distortion of a CI using linear system analysis tools. Instead of analysing the output of a modulated pulse train, we use the channel envelope to modulate a sine wave whose frequency is equal to the center frequency (CF) of the passband of that channel.

This allows us to carefully examine the effect of the frequency content of the envelope upon a pure tone. Since a user does not hear individual pulses, but they are merged for the most part into a continuous sensation resembling a tone, it makes sense to treat the pulse train as a tone. In addition, a square wave has a much more complex spectrum than a pure tone, and would greatly complicate the analysis. Third, we believe that the temporal information of the waveform has an effect on the frequency percept of the signal, and hence we want to know what that frequency is, and how it is affected by the preceding processing steps. Treating the envelope as a modulating signal, and an underlying sinusoid as a carrier, allows us to examine the effect of the modulating signal upon the output of the CI. Finally, even if one argues that a sine wave is not the proper representation of the carrier, but some alternate construct would do a better job of capturing the percept of an unmodulated pulse train, nevertheless, each frequency component of that construct will perforce be affected by the spectral content of the envelope in the same way as a single sinusoid would. By superposition, the output would be the sum of each of those modulated components. So, the most compact and atomistic representation, i.e., the most basic building block for analysis of such a system will still be a sinusoid, with any additional complexity being represented as a sum of additional modulated sinusoids. Therefore we can’t escape the need for analysis of a single sinusoid, no matter how we slice it. We further note, that some early work in simulating the sound of a CI in order to determine the minimum number of channels required for satisfactory intelligibility was performed by implementing such a sinusoid at the CF of each channel, hence it is an accepted method for recreating the auditory percept of a CI.[4]

We note one interesting point with the use of such a model. Because the envelope of a bandpass channel does not contain any absolute frequency information about the underlying signal, in effect, by modulating a sinusoid at the CF of the channel, we are incorporating place information into the model. The only way an accurate frequency percept could be produced by such a stimulus is if the place of the electrode holds some sway over the perceived frequency. But if we rely totally on place information, then any analysis we do of frequency content of the temporal characteristic of the signal becomes null and void, as all frequency content will default to the corresponding place/frequency relation for the particular location of the electrode. Hence, we are actually making use of a dichotomy in our analysis, and utilizing both spatial and temporal information in creating the actual percept. A ballpark percept is created by means of place information. But a more precise offset is produced by means of the temporal information of the envelope waveform. This may in some measure be analogous to the understanding we described in the Introduction, whereupon place information assists in producing better resolution than temporal information could produce, alone. In other words, they work together in combination to produce the final frequency percept. There is much to dissect here, in evaluating the appropriateness of this comparison, but we will leave it for future work. For now, we will proceed with the model, as described.

### 2.1 Modulation Theorem

In our earlier work,[1] we made heavy use of the modulation theorem which teaches that the frequency content of a modulated signal (CI output) has the same bandwidth as the modulating signal (channel envelope) centered about the frequency of the carrier (channel CF). Due to the fact that Fourier frequency components have both positive and negative frequencies, and for real signals, they mirror each other, the actual form of the output consists of two mirror-image sidebands extending above and below the carrier frequency. The designers of CIs, based on their psychophysical measurements of the frequency variations that a user can hear, decided to limit the cutoff frequency of the low-pass filter stage in the CI processor to approximately 300 Hz, depending on manufacturer. Whether or not one accepts their data and rationale, but the fact that this number is used in practice presents an iron ceiling on the maximum frequency excursions that can be produced by a CI above and below the CF of a channel. This means that if one has two adjacent channels, say with CFs of 6000 and 7000, respectively, there is absolutely no input stimulus that can produce the percept of 6500 Hz. The first channel is limited to excursions of 5700–6300 Hz, while the second is limited to 6700–7300 Hz, hence, the existence of a “dead zone” or spectral gap, in between. But the problem is actually worse than it appears.

### 2.2 Clarification on Spectral Gaps

At this point we stop to clarify a point that perhaps was not clearly explained in our earlier paper. The fact that there is never any output frequency between 6300 and 6700 Hz in our example does not imply that the user will hear silence with such an input. What does happen is that frequencies in the “dead zone” are transformed to frequencies in the “live zone”. The reason is that the nonlinear rectification stage has the capacity to generate new frequencies that were not present in the input. The modulation (multiplication stage) has the capacity to transform frequencies. Working together, they insure that all input frequencies do produce an audible output. But the modulation theorem guarantees that no output will be heard beyond the bandwidth of the input signal, which was limited by the low-pass filter. So without doing any complex nonlinear analysis, one can correctly say right off the top of his head that there will be a dead zone or “illegal frequency range” at the output. That is what we wrote early in our previous paper, based on simple reasoning that follows directly from the modulation theorem. But the more subtle point, which we only got to later in the paper, when we did more exact graphical analysis (we learned as we went along), is that there is no dead zone in the input. All input will actually be picked up by the system, but will be converted to “legal frequencies” i.e., within the live zone. Because of this confusion, readers actually emailed to say they tried sweeping across frequencies using various sound generators, but did not notice any actual dead zones—they heard everything. And we also received an email from Professor Louis Braida of MIT questioning their existence. I had the good fortune to later spend an hour with him drawing diagrams on his whiteboard, as we clarified this confusing point. There are no dead zones at the input of a CI, but there are dead zones at the output.

One can easily see how this will totally mess up the perception and enjoyment of music, in that certain notes on the scale will never be heard, but will be changed to other notes.

### 2.3 Frequency Transformation

Shortly we will demonstrate that the idea of legal frequencies, is actually too charitable. In fact, for single, pure tones, there really is only one frequency within the entire legal area that ever gets produced at the output. That is the CF of the channel. To see why, we need to work an example. This was done in great detail in our earlier paper, and exact, graphical, Fourier analysis was performed at every step as we traced the evolution of the signal from the input to the rectification to the low-pass filtering, to the modulation. And for each stage we graphed the results in both the time-domain and the frequency domain. Perhaps, the length and excruciating detail exhausted some readers, and caused them to skip over some of the material. So here we will give only a brief review, and then a simple arithmetic example to illustrate that the concepts are really not as difficult to work with as they may have appeared.

A few principles are in order. First, when one rectifies a signal, one adds in a DC (direct current)or zero-frequency component, because the original signal summed to zero, as positive and negative areas cancelled out. However, a rectified signal has a net positive area, producing a component at DC. Second, rectification also produces harmonics (integral multiples) of the input due to the sharp corners; or alternatively, in frequency domain thinking, due to it being a nonlinear operation. Third, the lowpass filter will remove any higher harmonics beyond its cutoff of 300 Hz. Fourth, when multiplying a DC value by a fixed frequency carrier (i.e., the CF of the channel, as in our chosen model), the output will be at the CF. If one multiplies a non-zero frequency, say a 100 Hz component of the envelope by the fixed frequency carrier at the CF, the output will be the fixed CF plus and minus 100 Hz. If the modulating frequency is 200 Hz, the output will be CF plus and minus 200 Hz, and so on. The lower frequency output is called the lower sideband, and the higher frequency output is called the upper sideband.

### 2.4 Example

Suppose we have a channel whose passband extends from 5,500 to 6,500 Hz, and with a CF of 6,000 Hz, as before. Further suppose that we play a pure tone sound of 5,600 Hz at the input. Let us trace the steps. The signal being within the passband of that channel, goes through the bandpass filter and reaches the rectifier. After rectification, it now has a component at 0 Hz, and also at 5,600 Hz and at 11,200 Hz and higher harmonics. It next reaches the low-pass filter. Only frequencies below 300 Hz are passed. The only component that makes it through is the DC component or 0 Hz. This component is then multiplied in the modulation stage by the fixed frequency carrier at CF, or 6,000 Hz. The result is an output of 6,000 Hz.

So, our 5,600 Hz output was transformed to 6,000 Hz. Not very nice. But perhaps understandable, since it was originally located in the illegal frequency range, and as we explained earlier, those get transformed to legal frequencies, and could not be heard otherwise, as there is no CI output in the illegal frequency range. This is illustrated in Figure 2.

**Figure 2.**
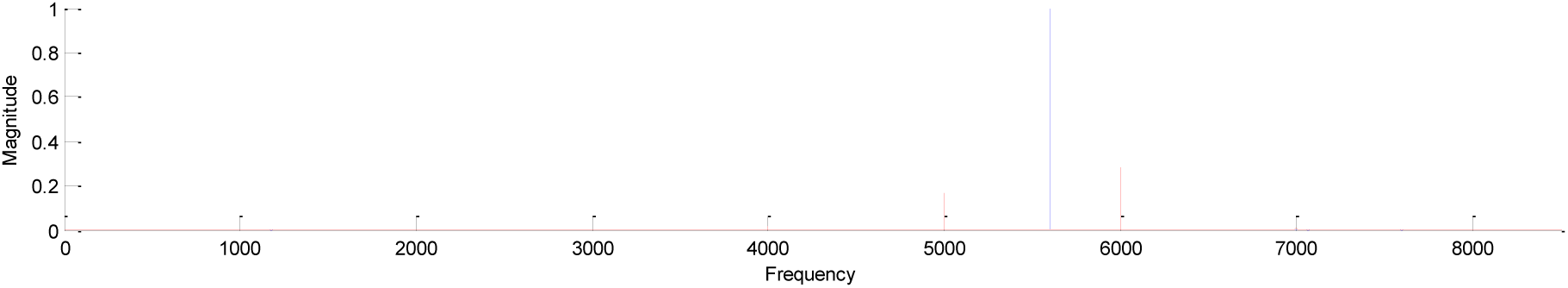
A single tone (blue) at 5,600 Hz, gets remapped (red) to center frequency of channel, 6,000 Hz, as described in text. (Note that due to properties of the 3rd order Butterworth bandpass filters used in this example, the signal was also picked up by the adjacent lower channel filter, centered at 5,000 Hz, and remapped to that frequency, as well. This double remapping, could conceivably be an additional source of distortion in CIs.)

But let’s now try another example where the signal was in the legal frequency range to begin with, and see what happens. We know that frequencies within 300 Hz or less of CF are legal, as we are entitled to as much bandwidth as our low-pass filter provides, which is 300 Hz. So let’s try an input of 6,200 Hz. Because the bandpass filter has a range of 5,500–6,500 Hz, the signal gets through nicely. It then gets rectified, producing components at 0 Hz, 6,200 Hz, 12,400 Hz, and higher harmonics. All frequencies above 300 Hz get blocked by the low-pass filter. The only one that makes it through is, again, the DC component at 0 Hz. After modulating the CF carrier with a 0 Hz component, the output is again at CF, or 6,000 Hz! This is illustrated in Figure 3.

**Figure 3.**
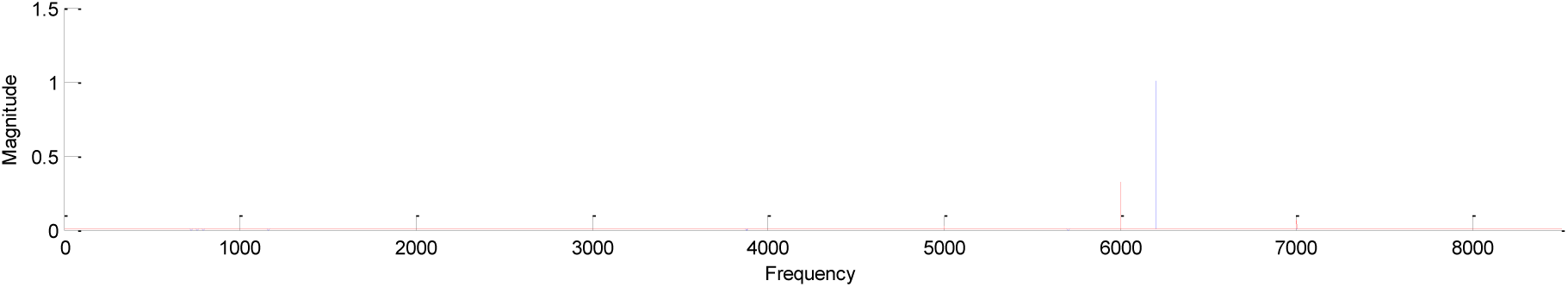
A single tone (blue) at 6,200 Hz, gets remapped (red) to center frequency of channel, 6,000 Hz, as described in text. (Note that due to the properties of the 3rd order Butterworth bandpass filters used in this example, the signal was also picked up weakly by the adjacent upper channel filter, centered at 7,000 Hz, and remapped to that frequency, as well. This double remapping, could conceivably be an additional problem of distortion in CIs.)

Very upsetting, as now, even when we started in the legal frequency zone, we still found that the input was transformed. Furthermore, both the input of 5,600 Hz and the input of 6,200 Hz, had identical outputs of 6,000Hz. You can’t tell the difference! This is what we referred to as a many-to-one transformation in the Introduction. Multiple inputs produce an identical output. This makes music even more difficult to understand than we would have thought before. And this is why even the legal frequency zone is almost always quiet in the case of pure tones, except for one single frequency…the CF of the channel!

Suppose we would plot all the possible single-tone frequency inputs, and their possible outputs. It should seem plausible that over most of the range, we would get a staircase function. All frequencies in one passband get transformed to that channel’s CF. Then, as we leave that passband and enter the next, all frequencies now get transformed to the CF of the second channel, and so on.

These rules, which we confirmed with exact, graphical Fourier analysis in our earlier paper, can be consolidated into the following plot shown in Figure 4. We arbitrarily chose the following list of frequencies to be the CFs of the channels. We used easy, round numbers for clarity, but note that in the real world, the band centers are nowhere near as simple, and can be related through any of a number of schemes which do not yield integral or evenly spaced values.

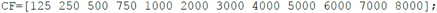

**Figure 4.**
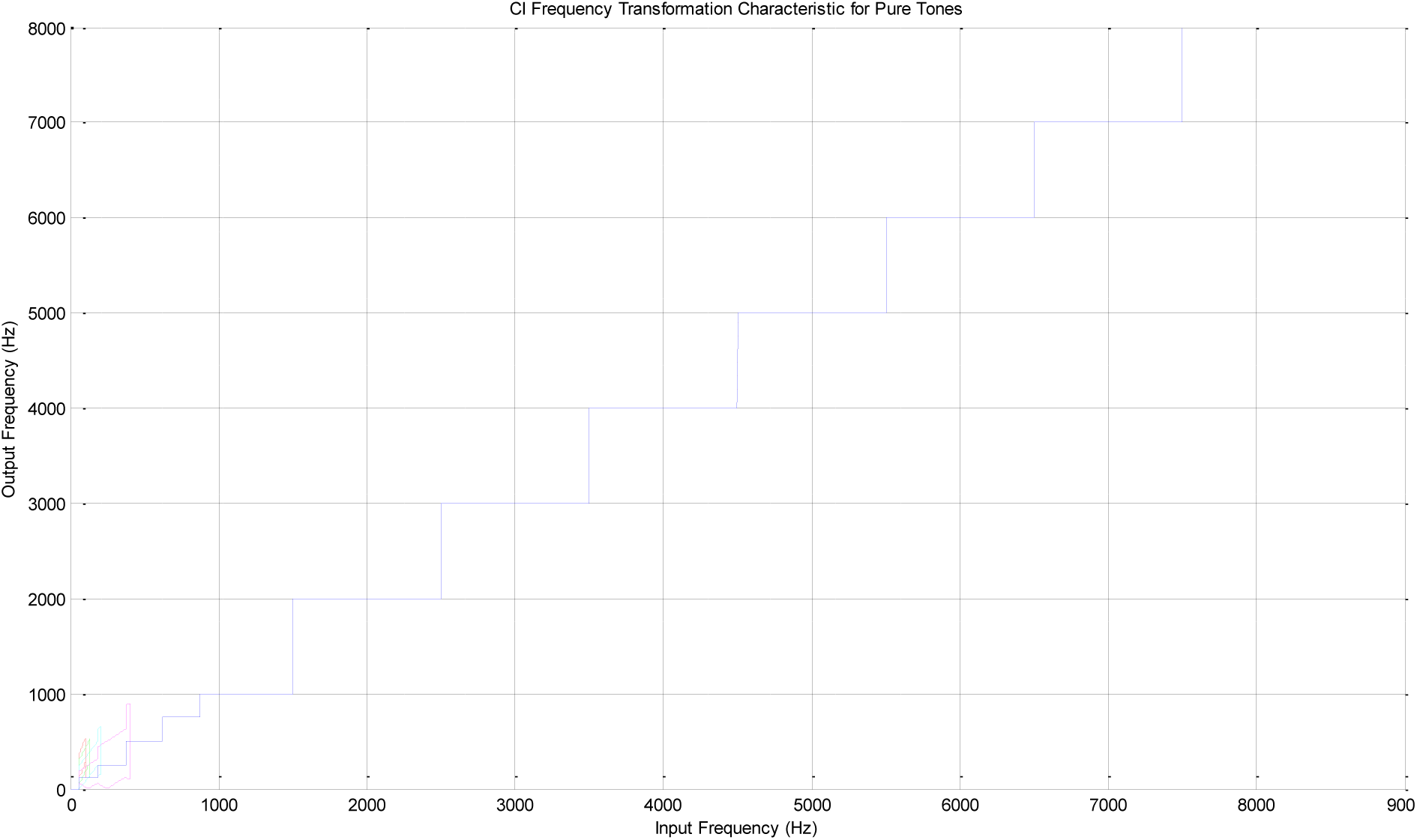
A plot of input vs. output frequency through a CI for pure tones, using band center frequencies as listed in text. For very low frequencies, some sidebands are able to make it through the lowpass filter (with cutoff here at 400 Hz), and these are indicated as additional pairs of colored lines above and below the main blue line which tracks the fundamental frequency output. Note that negative frequencies get converted to positive values, and hence the jagged lines at bottom. A replot without such conversion is provided in next figure, for clarity.

**Figure 5.**
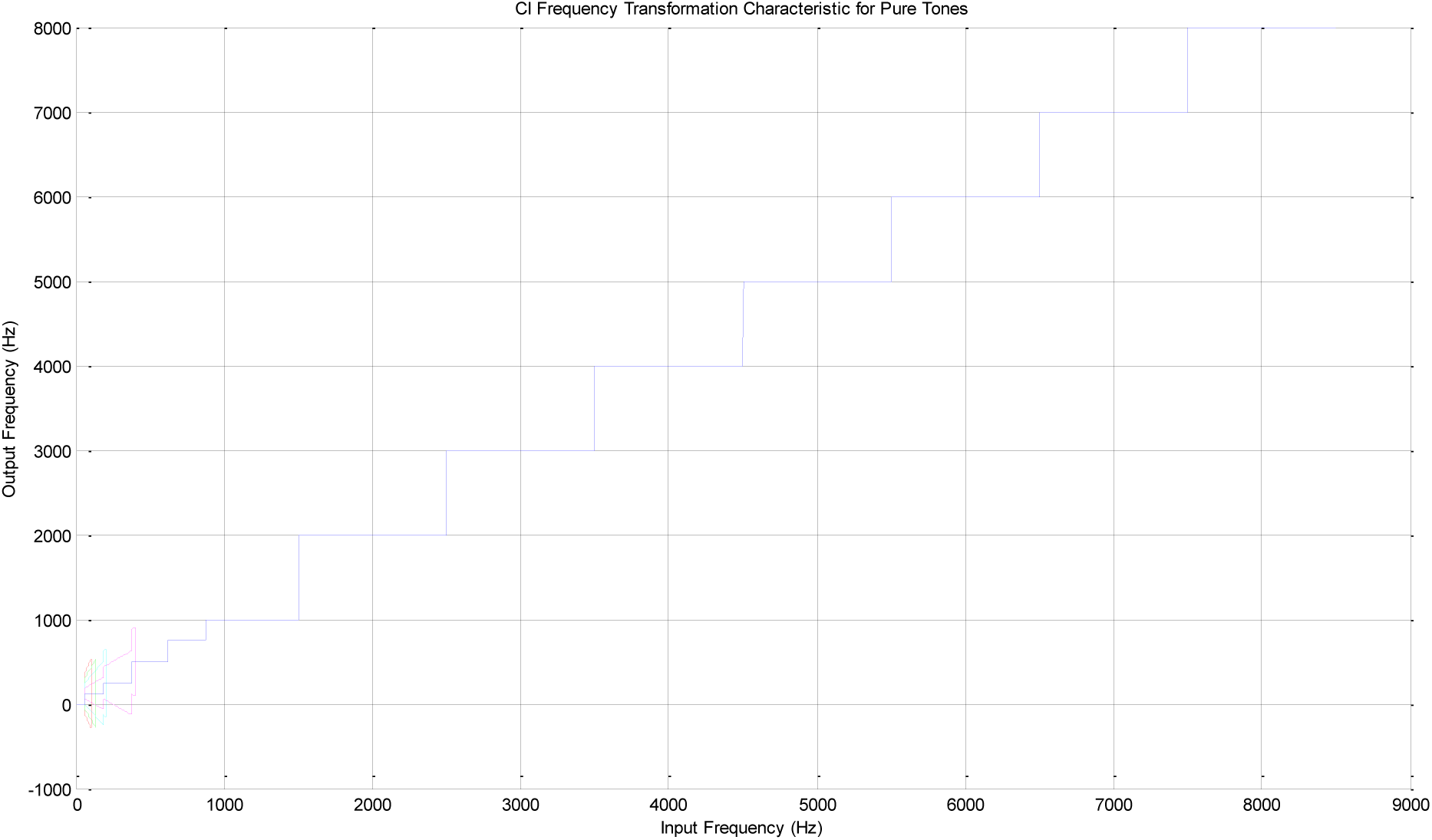
Same plot as previous, except allowing frequencies to go negative for illustrative purposes, to better see the trajectory of the sidebands (various color pairs) above and below the fundamental (blue).

The output frequency is a staircase function of the input frequency, as we reasoned. We emphasize that this holds only for pure tones. We also note that at the very low end of the spectrum, certain sidebands do make it past the low-pass filter, and a single input can have multiple output components. These will track linearly with the input, until they exceed the cutoff frequency of the lowpass filter. Hence, they have a diagonal, rather than staircase appearance. We show the first upper and lower sidebands as a pair of lines of one color, and the second set of sidebands, as a pair of a different color, and so on. We note that this plot was not made using actual Fourier analysis, but is an illustration of what we expect according to the rules of such analysis. Later, we will use exact examples, where we constructed a simulated rectifier, lowpass filter, and modulator, and actually did compute a graphical Fourier analysis of the input vs. output for certain representative cases.

### 2.5 Complex Inputs

The rules we have laid out hold only for single, pure tones. As we showed in our earlier work, when using complex tones, a number of complications may arise. These can be due either to mixing of components within a single channel, or interactions between components of separate channels. The first has the ability to actually create new frequencies that were not present at the input, while the second can cause dissonance between components of different channels that are out of tune with each other.

#### 2.5.1 Out of Band Interference

We begin with the second case, as it is easier to understand. As we saw, single tones within a bandpass filter get transformed to the CF of that filter, according to our model. But complex tones like vowels and musical notes have multiple harmonics that are all integrally related to each other, and whose aggregate produces a nice pleasing periodic signal, since all the components repeat within the same cycle. However, if a certain harmonic falls within the passband of one channel, and another harmonic falls within the passband of a different channel, then the outputs of the channels may no longer be harmonically related (as integral multiples of a common frequency). There is no reason to assume that the CFs will have any simple relationship to each other. Harmonics of a single instrument may thus become mistuned with respect to each other. Instead of having a series of tones at exactly 100, 200, 300, 400… Hz, the output might be 103, 198, 315, 407… Hz, etc. These will cause much grating and dissonance when played together. A human voice will also sound poor under such conditions.

#### 2.5.2 Inband Interference—Intermodulation Distortion

The situation becomes even more complex when two tones interact within a single channel. As one progresses in frequency, the bands generally become wider, and can admit multiple harmonics from the same instrument. This can happen in the lower channels, as well, in certain cases. Whenever there are multiples inputs to a nonlinear system, the result is often unpredictable, and as we saw in our earlier work, new frequency components can be created that were not present in the input, and are greater in number than the number of input frequencies. In that paper, we showed an example within a single channel, and left it at that, and speculated that this would cause severe distortion as it is likely to occur in multiple bands and these would all interact dissonantly. In that work, we made no effort to explain the relationship between the input and output frequencies for multiple input cases; we simply recorded what the Fourier analysis showed. In general, nonlinear systems are like the wild west, each different than the next and one doesn’t usually expect to find too many unifying principles like there exist for linear systems.

However, in our fruitful discussion with Professor Lou Braida, he suggested that using the rules of intermodulation distortion actually gives a fairly accurate way of understanding the expected outputs from our model. These rules are that integral multiples of each input frequency can be generated, but that each of these frequency components can be added and subtracted from each other. Some will be out of the range of the lowpass filter, but some will lie within it, and can proceed to the output.

As an example, if one uses inputs of 5800 and 5900 Hz, using the channel configurations as in our previous example, there can be a difference frequency generated of 100 Hz, in addition to the two input frequencies and their harmonics. And there can be a difference frequency between the two second harmonics producing a 200 Hz signal. These can get through the lowpass filter of 300 Hz cutoff. Now when these modulate the carrier frequency, we produce upper and lower sidebands of 6,000 +/- 100 Hz and 6,000 +/- 200 Hz. These will yield components of 5,800, 5,900, 6,000, 6,100 and 6,200 Hz. So when we have multiple input signals, the output is not always remapped straight to the single frequency at the CF of the band, as it is for single, pure-tone inputs. Similarly, in cases of wideband noise. In all these cases, there will be multiple components within the legal frequency range of the system. We stress that never will any components be produced in the illegal frequency range, as that is an iron-clad rule set by the characteristics of the lowpass filter.

While in this simple example, it may seem that some of the output components turned out to be at the same frequencies as the input components, that was a fortunate coincidence, since we used round numbers for the channel CFs and for the input frequencies, which matched nicely. We will shortly see a graphical simulation where this does not occur, as we change the fundamental frequency from 100 Hz to 150 Hz.

It turns out, therefore, that there are subsets—particularly well-characterized nonlinear systems—for which simple integer rules hold in computing the frequencies of distortion products. These include rectifiers and certain other common circuit operations.

### 2.6 Complex-Tone Example

The following plots, Figure 6 and Figure 7, show a complex tone, which could be a vowel or musical note consisting of a 150 Hz fundamental, along with all of its harmonics plotted in blue, and the corresponding output plotted in red. Third order Butterworth filters were used for the bandpass filters, and a fifth order Butterworth filter for the lowpass filter. The lowpass cutoff frequency was set at 300 Hz.

**Figure 6.**
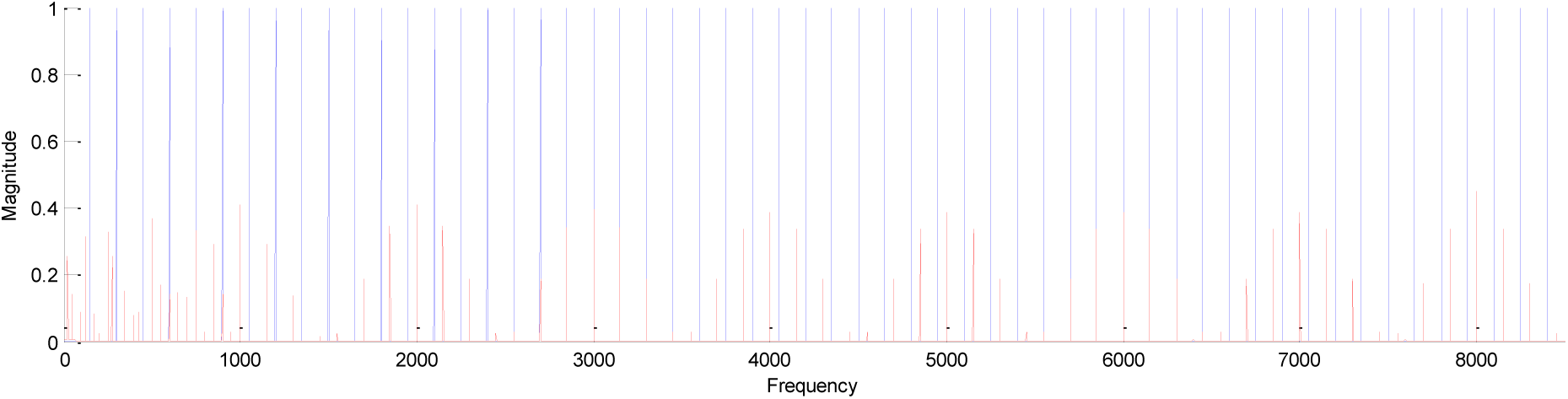
The complete response (red) to a single complex tone (blue) consisting of a fundamental at 150 Hz, and all its harmonics. Note that when multiple input tones fall within the pass band of a single channel, it is possible to generate a response away from the CF of channel in the legal frequencies, but never within the dead zone or illegal frequencies in between channels which are always verboten.

**Figure 7.**
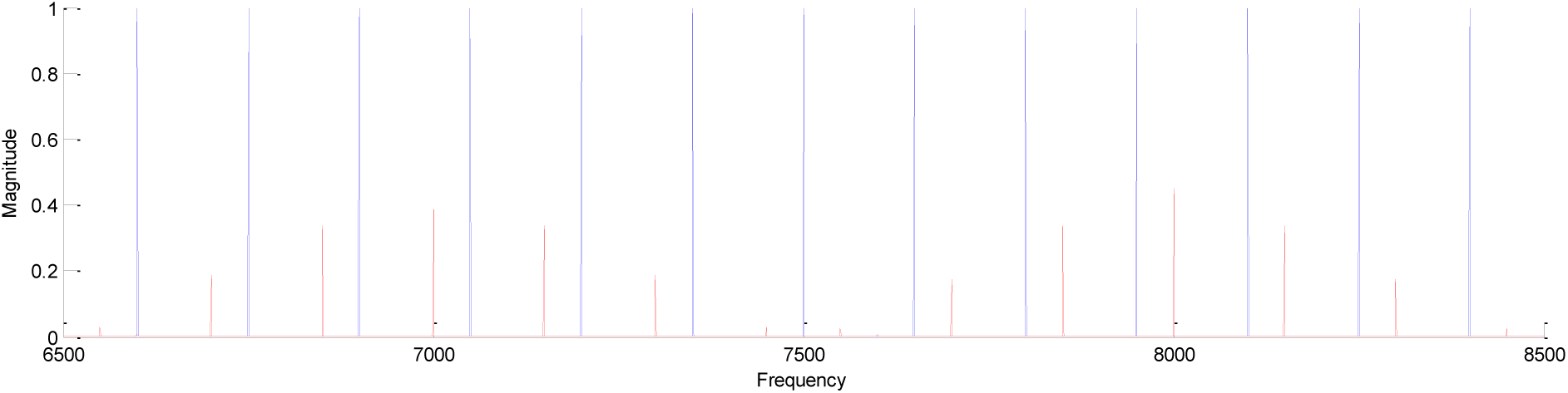
Here we zoom in on previous response in order focus on the region between 6500 and 8500 Hz, comprising the highest two channels. Note that the spacing between harmonics of the original (blue) is 150 Hz throughout. The spacing between the harmonics of the response (red) is also 150 Hz, but origin resets at each channel CF. At the CF of first channel, 7,000 Hz, there is a response, and then additional response components (sidebands) above and below CF, separated by 150 Hz. The same occurs for the channel with CF of 8,000 Hz. But neither of these are consistent with the definition of a harmonic set, as they are offset from being true multiples of a single fundamental, and hence would yield an aperiodic tone. They are not consistent with each other, either, as for the 7,000 Hz group, they are consistently 50 Hz below the correct value, while for the 8,000 Hz group, they are consistently 50 Hz above the correct value. Hence they would clash with each other, as well. Note that in real life, the CFs are not set to round numbers like we have used here, but can be logarithmically or otherwise related, making the situation even more complex.

The first feature which is noticeable is that the responses cluster around the CFs of the channels. This is expected from our analysis, in which we showed that there must be a falloff in response as one moves away from the CF, due to the action of the lowpass filter. The second feature is that the components of the original signal are regularly spaced throughout at exactly 150 Hz intervals, as would be expected from a harmonic set. The output components are also spaced at 150 Hz intervals, but these are offset from the CF of each band. They are due to the rectification process producing harmonics of 150 Hz, and sum and difference frequencies of these harmonics, which are also multiples of 150 Hz. These become mapped as upper and lower sidebands of each channel CF. However, they do not fit the definition of a harmonic set, since while they are separated by 150 Hz, they are not common multiples of 150 Hz. Furthermore, the sidebands which are offset from one channel’s CF, may be out of sync with the sidebands that are offset from the next channel’s CF. This can be seen in Figure 7, where the red lines are 50 Hz lower than the blue lines in the left side of the plot where the channel CF is 7,000, but become 50 Hz higher than the corresponding blue lines in the right side, where the channel CF is now 8,000 Hz. This will produce dissonance, because the summed output will be aperiodic. The frequencies are not integral multiples of each other, and hence don’t begin and complete a cycle at the same time.

One final observation is that for low input frequencies, there is a possibility that two or more higher-order distortion components from a single input harmonic can pass through the low-pass filter. This produces additional tightly spaced spectral lines in the low frequency range, compared to the more widely separated output components in the higher frequency range.

### 2.7 Multiple Talkers

The final example we will examine shown in Figure 8 and Figure 9 is the case of two complex tones, such as when two speakers are talking at the same time. The first speaker has a fundamental of 149 Hz, and the second, 150 Hz. In addition to the distortion produced as in the previous case of a single speaker, there are now intermodulation components between the frequency components of each speaker. The harmonics themselves each map to sidebands offset +/- 149 and +/- 150 from CF of 8,000 Hz, and also have second-order distortion products of +/- 298 and +/- 300. In addition, we have weaker intermodulation components F_1_-F_2_ and 2F_1_–2F_2_ generating frequency offsets of +/- 1 Hz and +/- 2 Hz and so forth from CF, as can be seen in Figure 9

**Figure 8.**
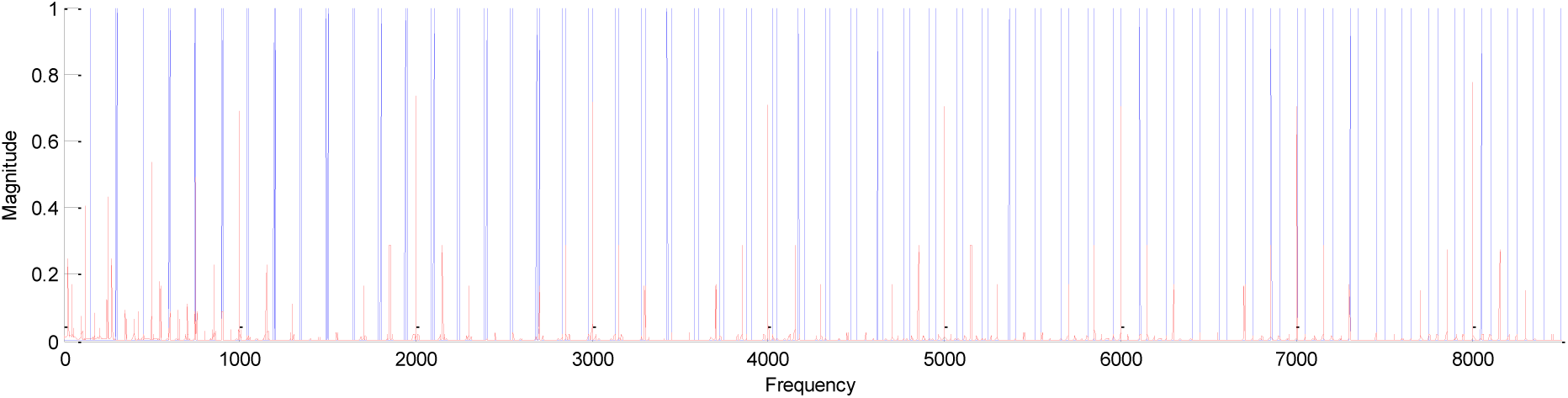
The complete response (red) to a set of two complex tones (blue). The tones had fundamental frequencies of 149 and 150 Hz, respectively. Each consisted of a fundamental and all integral harmonics up to 8,500 Hz. The response is always clustered around a channel CF, and falls off with the characteristics of the lowpass filter used in the CI processor following rectification.

**Figure 9.**
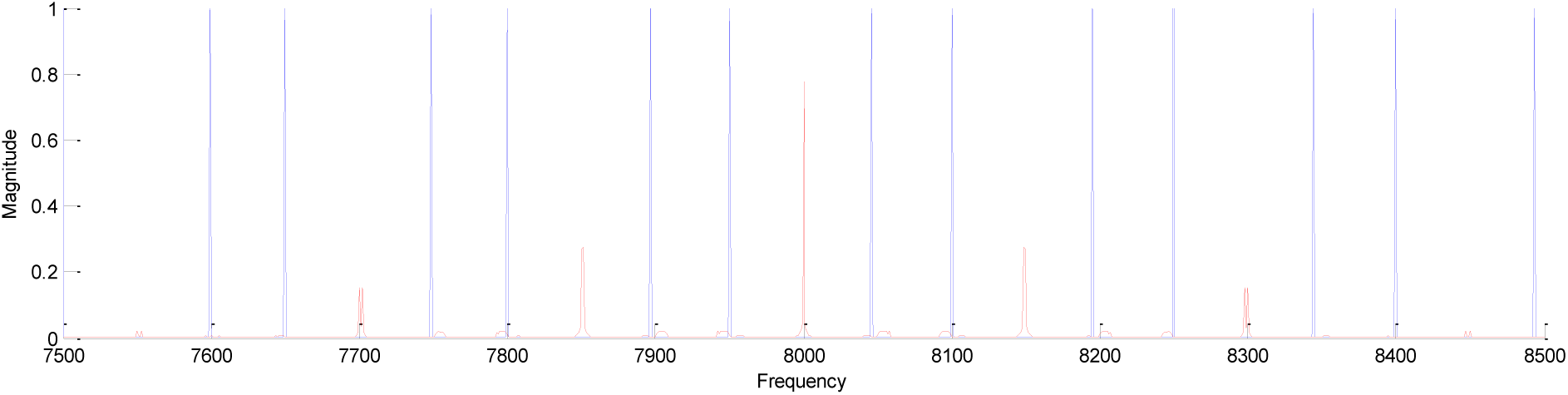
A zoomed view of previous figure centered about the highest channel CF at 8,000 Hz. Note that neither of the original (blue) series has any component at 8,000 Hz, while response (red) is mapped to 8,000 Hz, the CF of channel. Also note that components of the original two series (blue) become farther apart from each other as frequency increases, but the responses (red) to each series coincide at 8,000 Hz, are 1 Hz apart at 8,149 and 8,150 Hz (first upper-sideband harmonics) and also at 7,850 and 7,851 Hz (first lower-sideband harmonics), then are 2 Hz apart at 8,298 and 8,300 Hz (second upper-sideband harmonics) and also at 7,700 and 7,702 Hz (second lower-sideband harmonics). These incorrect and closely spaced components will lead to a beating or dissonant effect that gives rise to a buzzlike, raspy or grating sound quality, in addition to being out of tune, musically.

The net effect is that at 8,000 Hz, we have an incorrect component due to the DC component from each harmonic which maps directly to CF. Then we have doublets at +/- 149 and +/- 150, and also at +/- 298 and +/- 300 from the CF. These doublets will certainly create a very unpleasant sensation, as their components are right near each other. No such closely-spaced components exist in the original signal in the higher frequencies. So the CI output deviates greatly from the original. This may explain the great difficulty CI users have in noise.

## 3 Discussion

We have demonstrated that if we accept our model, which attempts to apply traditional signal processing techniques to the analysis of the spectral components of a signal as it traverses each stage of CI processing, much distortion is generated.

This manifests itself in a number of different ways:

- Spectral gaps or dead zones in frequency response
- Frequency transformations
- Staircase like response to smooth frequency changes
- Many-to-one response producing same output for different tones—ambiguity
- Generation of nonexistent components and loss of true components
- Dissonance

We hear these problems very clearly, and they greatly impact our ability to hear voices, even in quiet, but especially in noise. It can be hard even to distinguish a male voice from a female voice, since they both sound like they have a bad sore throat.

For music, the situation is even worse, as melodies become completely unrecognizable, as correct notes are not heard, and often two different notes sound the same. We recently received correspondence regarding another user who made the exact same observation. Notes seem the same, as you increase frequency, until you hear a sudden jump to a new frequency. We believe this clearly describes the staircase effect we described and diagrammed in the previous section.

We note that it may be possible to detect note changes, even if the fundamental is in a flat part of the staircase curve, because perhaps it may have harmonics that fall near a border, and may jump into the next bandpass channel while the fundamental is still in its original channel. Nevertheless, this is a very poor and inexact method, which has limited utility, and is very un-natural, in that normal hearing listeners hear all harmonics moving together.

We have allowed ourselves throughout to use the temporal features of waveforms to characterize the frequency percepts that would be heard. We acknowledge that some researchers may reject our model out of hand, because they believe that spatial cues play the major role in frequency perception, and hence temporal analysis is irrelevant. We strongly object, since if that were true, a listener would not be able to hear beating between two closely spaced frequencies. But we hear it clearly, and believe that all users would notice these fluctuations, although one could conduct a formal study to verify. If that is the case, one cannot escape the fact that temporal interactions do play a role for CI patients. We are also very bothered by the fact that by using a low-pass filter of a few hundred Hz, designers may be creating a self-fulfilling prophecy that users cannot distinguish frequencies well, so let’s take it away from them. We do not believe that whatever psychophysical tests were used can justify this decision.

Furthermore, if our analysis is correct, then the source of the most prominent distortion is due to CI processing algorithms, and is not biological in origin. This eliminates interelectrode interference or current leakage as a cause, via Occam’s razor. I.e., if one can show that a certain cause produces a certain effect, there is no need to suspect that there may be a second cause which has not been detected.

Consequently, we don’t believe there is justification for limiting the number of channels in order to keep their electrodes as far apart as possible. And we also do not believe that interleaved pulse stimulation is necessary. One could use continuous stimulation. And one could also eliminate all the envelope processing steps, and simply pass the original bandpass signals, with suitable amplitude adjustments, so as to be compatible with cochlear signal levels, to each electrode.

We believe that CIS introduces additional additive distortion of its own in the form of switching noise, which is heard as a continuous crackling and hissing on top of the output. We read an online report of a patient complaining to Dr. Don Eddington about this noise, but again, not being an actual user (although he is one of the world experts and helped ne immensely), he searched and could not find any malfunction. Only a user who hears the switching noise would realize that it is part of the normal operation of the device. We have confirmed this with a user of another brand, as well.

## 4 Conclusion

Based on this work and our earlier work, we believe that better performance could be achieved by eliminating envelope processing and using the unadulterated signal. Envelope processing does not simply eliminate certain potentially useful information. It actually rewrites the frequency content of the signal. We do not believe researchers have stopped to examine how severe this effect is. Speech can be heard, because it is extremely robust, and even four bands of noise can produce intelligible speech. But in difficult situations, it becomes extremely hard to decipher. Even in the best case, the distortion is extremely irritating. It sounds buzz-like, raspy and grating, as if the speaker had a sore throat. Music is much worse, as we described.

While older studies seemed to show that certain patients did better with CIS processing, however, the number of channels and the quality of electrodes has gone up since then. We believe that with the Med-El unit we do better than users of other manufacturers due in large part to their extremely long electrode which provides good low frequency coverage.

Because we don’t believe that interelectrode interference is a concern, it should be possible to add many more channels, up to as many as practical engineering considerations will allow, such as space requirements for wires, adequate transmission through skin, and power consumption, etc. The interference which patients have reported is likely due to internal generation of distortion products, which appear to come from other channels, not actual current leakage from those channels.

We do have a suggestion for how to improve the bandpass filters in continuous-time systems so that better intelligibility can be achieved. This is based on certain work we did as part of our thesis research, where we showed that exponential filters have certain unique properties that may mimic the responses of natural cochlear filters. We will leave this for a future paper.

But we are optimistic that because of the excellent loudness and ability to hear in many situations, that the underlying technology will continue to improve. We hope that manufacturers and researchers will consider the ideas and analysis expressed here with an open mind, despite that they may be heavily invested in alternate schemes and ways of thinking.

